# Engineering bacteriocin-mediated resistance against plant pathogenic bacteria in plants

**DOI:** 10.1101/649178

**Authors:** William M. Rooney, Rhys Grinter, Annapaula Correia, Julian Parkhill, Daniel Walker, Joel J. Milner

## Abstract

*Pseudomonas syringae* (*Ps*) and related plant pathogenic bacteria are responsible for losses in diverse crops such as tomato, kiwifruit, pepper, olive and soybean. Current solutions, involving the use of chemicals and the introduction of resistance genes, have enjoyed only limited success and may have adverse environmental impacts. Consequently, there is a pressing need to develop alternative technologies to address the problem of bacterial disease in crops. An alternative strategy is to utilise the narrow spectrum protein antibiotics (bacteriocins) used by diverse bacteria for competition against closely related species. Here, we demonstrate that active putidacin L1 (PL1) can be expressed at high levels *in planta* and expression of PL1 provides effective resistance against diverse pathovars of *Ps.* Furthermore, we found that strains which evolve to become insensitive to PL1; lose their O-antigen, exhibit reduced motility and are less virulent in PL1 transgenic plants. Our results provide proof-of-principle that transgene-mediated expression of a bacteriocin *in planta* is an effective strategy for providing disease resistance against bacterial pathogens. Genetically modified (GM) crops expressing insecticidal proteins have proved extremely successful as a strategy for pest management; expressing bacteriocins to control bacterial disease may have a similar potential. Crucially, nearly all genera of bacteria, including many plant pathogenic species, produce bacteriocins, providing an extensive source of these antimicrobial agents.

**SIGNIFICANCE:** With the global population to surpass 9 billion by 2050 there is a huge demand to make industrial farming as efficient as possible. A disadvantage of industrial farming is the lack of genetic diversity within crop monocultures, which make them highly susceptible to diseases caused by plant pathogenic bacteria like *Pseudomonas syringae*. Bacteriocins are narrow spectrum protein antibiotics which are produced by all major bacterial lineages. Their main purpose is to eliminate competitor strains to establish dominance within a niche. By arming plants with bacteriocins we can increase the genetic toolbox used to engineer crops to be resistant to specific bacterial plant pathogens.

---

*Pseudomonas syringae* (*Ps*) is a Gram-negative bacterial plant pathogen. The *Ps* species complex consists of over 50 known pathovars (pv.) all of which cause different diseases such as bacterial speck, spot and blight disease on tomato, beans, tobacco and a large number of agronomically important crops (1). Once a plant pathogen is introduced into a crop it can spread rapidly due to the lack of genetic diversity in the commercial crop varieties sown (2). A recent example of this is the pandemic caused by *Ps* pv. *actinidiae* (Psa), which is currently causing massive damage to the global kiwi fruit industry.

The emergence of canker disease on commercial kiwifruit (*Actinidia spp*.) varieties has been well documented since the early years of *A. deliciosa* domestication in Japan in 1984 (3). Since then Psa has been detected in China (1984), Korea (1988) (4) and Italy (1992) (5). In 2008, a hypervirulent strain of Psa was isolated in Italy on *A. chinensis* and was subsequently found in other neighbouring European countries, South America and Asia (6, 7). These aggressive forms of Psa caused massive economic devastation to kiwi growing countries; for example Psa was detected in 37% of New Zealand’s kiwi orchards (8).

Current solutions, involving the use of chemicals have enjoyed only limited success, encourage the evolution of resistance among the bacterial populations and may have adverse environmental impacts (9, 10). Furthermore, the introduction of resistance genes like EFR and Bs2 into tomato have been shown to be successful at providing resistance against *Ps.* However, there is a distinct lack of diversity of genes identified that can be introduced into commercial crops (11). Therefore, there is a pressing need to develop new technologies to introduce disease resistance against economically important plant pathogens like *Ps*.

The large *Ps* species complex promotes intense selective pressure on individual species to evolve mechanisms to eliminate inter- and intra-species competition in their environmental niche. One mechanism is the production of bacteriocins, which are narrow spectrum proteinaceous antibiotics that target and kill related bacterial species. Utilisation of the highly targeted antibiotic activity of bacteriocins provides a potential route to crop protection against specific bacterial pathogens with minimal impact on the wider microbial community (12).

A number of prospective bacteriocins have been identified in *Pseudomonas spp.* including putidacin L1 (PL1), a 30 kDa lectin-like bacteriocin that has been shown to be very potent against *Ps* pv. *syringae, lachrymans* and *morsprunorum* (13, 14). The lectin-like bacteriocins bind to D-rhamnose containing oligosaccharides incorporated into lipopolysaccharide (LPS) on the bacterial surface (15, 16). This facilitates the docking of PL1 to the cell surface and interaction with the outer membrane insertase BamA, leading to death of the cell via an unknown mechanism (17). Recent interest in producing and using bacteriocins for the treatment of bacterial infections in humans have promoted attempts to biopharm in plants bacteriocins with activities against *E. coli*, *Salmonella* and *Pseudomonas aeruginosa* (18–20). The successful demonstration that active bacteriocins can be expressed in plants suggests that PL1 could also be expressed *in planta* in an active form, thereby protecting plants against *Ps* infection.

In this study, we conclude that transgenic expression of a bacteriocin in planta can provide robust disease resistance against the bacterial phytopathogen *Ps*. We demonstrate that active PL1 can be efficiently expressed in both *Nicotiana benthamiana* (*N. benthamiana*) and Arabidopsis. In addition, transient expression in *N. benthamiana* and stable expression in Arabidopsis can provide quantitative and qualitative disease resistance against PL1-sensitive strains of *Ps*. Moreover, we show that mutations associated with PL1-insensitivity were linked with a key lipopolysaccharide biosynthesis machinery and that these mutations had a fitness cost in PL1-expressing plants.

## RESULTS

### PL1 has a narrow killing spectrum

To determine the killing spectrum of PL1 against *Ps* pathovars, recombinant PL1-His_6_ produced in *E. coli* and purified protein were used to assess killing activity against a panel of 22 diverse *Ps* pathovars. These include pathogens of kiwifruit, locust bean, oat, soybean, cucumber, cabbage, cherry, plum, olive, pear, maize, lilac and tomato. Of the 22 strains tested, 10 (from 6 different *Ps* pathovars) were sensitive to PL1 (Table S1 and Fig. S1) including all 3 members of the *syringae* group. All 4 members of the tomato group were resistant to PL1. In conclusion, PL1 has a very specific killing spectrum which makes it an ideal candidate to express in plants.

### PL1 provides robust disease resistance against *Ps* pv. *syringae* LMG5084 in *N. benthamiana*

Previously, bacteriocins active against human pathogens have been expressed in *N. benthamiana* leaves and leafy green vegetables (18–20). To express PL1 *in planta*, a construct encoding PL1 with an N-terminal 4 × c-Myc tag was cloned into a Ti binary vector and transiently expressed in leaves of *N. benthamiana* using agroinfiltration. By 3 days post-infiltration, leaf extracts showed high levels of PL1 in western blots and in spot tests using the PL1-sensitive strains. PL1 expression correlated with a killing ability equivalent to 0.35% of total plant protein (~5 µM), demonstrating that active PL1 can be produced efficiently in leaves (Fig. 1a,b).

**Figure 1.**
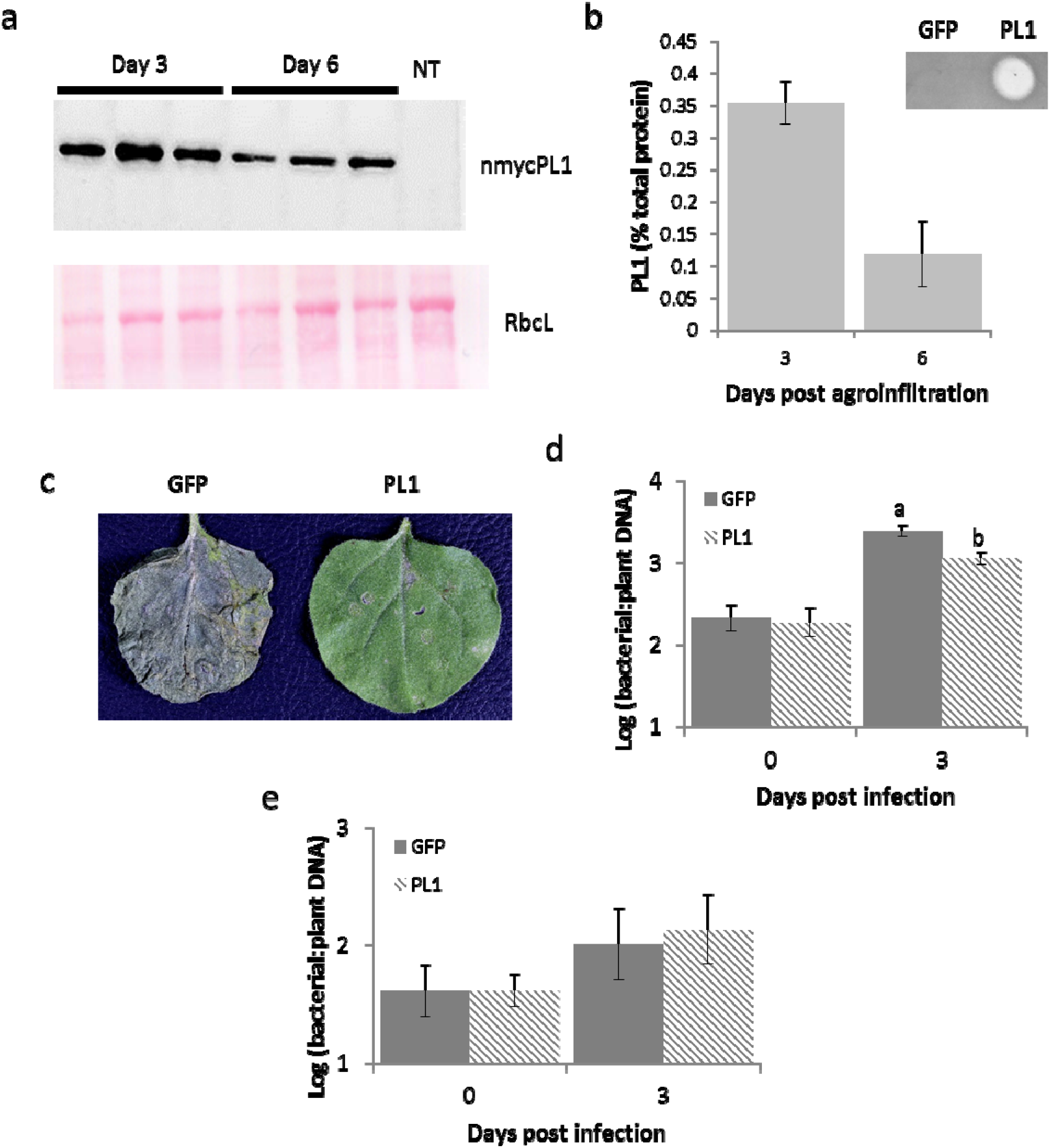
Transient expression of PL1 in *N. benthamiana* attenuates the growth of PL1-susceptible strains but not PL1 resistant strains. **a**, PL1 can be expressed at high levels in *N. benthamiana*. Western blot of myc tagged PL1 expressed in *N. benthamiana* leaves 3- and 6-days post infiltration with the corresponding Ponceau S staining with a non-transgenic control (NT). **b**, % Total protein and activity of PL1 from transient expression. Serial dilutions of whole protein extracts from *N. benthamiana* leaves 3- and 6-days post infiltration were spotted onto lawns of LMG5084 to estimate the percentage of PL1. A representative spot test with a GFP control is shown. Error bars represent the standard deviation of 3 independent replicates. The above panel represents a lawn of LMG5084 with spots of *N. benthamiana* extracts expressing either PL1 or GFP (control). **c**, PL1-expressing leaves showed qualitative resistance to LMG5084. *N. Benthamiana* leaves either expressing PL1 or GFP were infiltrated with LMG5084 (3 days post agroinfiltration) and left for 7 days post infection for symptoms to develop. *N. benthamiana* leaves 3 days post agroinfiltration expressing either GFP or PL1 were syringe infiltrated with **d**, PL1-expressing leaves show quantitative resistance to LMG5084 but not DC3000. LMG5084 and **e**, DC3000. Genomic DNA was extracted from the leaves expressing GFP or PL1, 0 and 3 days post infection and the bacterial load was measured by qPCR relative to plant tissue. Error bars represent standard error of 3 independent replicates. Statistical significance within the same time points was revealed using a one-way ANOVA and post hoc Tukey’s T-test. Letters denote statistically significant groups (p< 0.05).

After establishing that PL1 can be expressed transiently at high levels in leaves, we challenged these leaves with *Ps* to establish whether we could see a qualitative difference in disease symptoms. Three days post agroinfiltration (now denoted as day 0), the leaves were inoculated with LMG5084 (a pathovar that is highly sensitive to PL1 –Table S1) or a PL1-insensitive strain DC3000. Leaves were observed for symptom development and bacterial growth was measured over the subsequent 3 days. Infiltrating leaves with *Agrobacterium* has been shown to induce immune responses that inhibit the growth of *Ps* in subsequent inoculation (21–23). We therefore compared the growth of *Ps* in leaves transiently expressing PL1 following agroinfiltration with a control of leaves transiently expressing Green fluorescent protein (GFP), a non-bactericidal protein. With LMG5084, leaves expressing PL1 showed striking reductions in symptom severity (mild chlorosis only) compared to controls expressing GFP which exhibited black mottling and extensive necrosis by 7 days post-infection (dpi) (Fig. 1c).

When we measured bacterial load in leaves expressing PL1*, Ps* titres were 5-log units lower than in non-agroinfiltrated control leaves and crucially 3-log units lower than in leaves expressing GFP (Fig. S3a). We also showed that the process of syringe infiltration with buffer does not affect the growth of *Ps* and that GFP expression did not further affect the growth of *Ps* compared to the empty vector control (Fig. S2). The PL1-resistant strain DC3000 showed no difference between titres of *Ps* in leaves expressing PL1 or GFP (Fig. S3b). Titres of LMG5084 (but not DC3000) recovered from leaves immediately following inoculation (i.e. 0 dpi) were unexpectedly ~2-log units lower in leaves expressing PL1 than in leaves expressing GFP or non-infiltrated controls (Fig S3a,b). We suspected that this was a result of killing post-extraction following release of PL1 during grinding and confirmed this by mixing leaf extracts from PL1-expressing leaves with leaf extracts from infected non-expressing leaves (Fig. S4a,b). We therefore developed an alternative assay using qPCR to measure the quantity of bacterial genomic DNA relative to plant DNA in leaf extracts (24).

A standard curve showed a linear relationship between bacterial titres (colony forming units) and bacterial DNA recovered *in planta* (Fig. S5). Next, we repeated the infection of leaves expressing PL1 and GFP for both LMG5084 and DC3000. By 3 dpi we observed significantly reduced levels of bacterial DNA (p-value=0.031) in PL1-compared to GFP-expressing leaves following inoculation with LMG5084 but not DC3000, consistent with the direct measure of bacterial titres (Fig.1 d,e). DNA levels do not distinguish between living and dead bacteria and therefore overestimate the titres of living bacteria within the leaf and underestimate any differences. In conclusion, we showed that PL1 can be expressed to a high level in *N. benthamiana* leaves and that the expression of PL1 correlates with both qualitative and quantitative disease resistance against the PL1-sensitive strain but not the insensitive strain DC3000.

### PL1 provides robust disease resistance against *P. syringae* pv. *syringae* LMG5084 in Arabidopsis

To better understand the efficacy of PL1-mediated resistance, non-transgenic (NT) Arabidopsis plants were transformed to express c-myc PL1 and we selected homozygous PL1-expressing transgenic lines. We characterized 3 lines, PL1(1-2), PL1(2-1) and PL1(6-1) which had varying levels of PL1 expression from 0.13-0.67% or total protein extract (1.5 −10 µM) (Fig 2. a,b). The lines were infected by spraying them with suspensions of the PL1-susceptible pathovar LMG5084. As a control, the PL1-resistant line DC3000 was used. Bacterial titres we measured over 3 days. Titres of LMG5084 were significantly lower in PL1(1-2) and PL1(2-1) with p<0.001 and p< 0.004, respectively. The greatest reduction in growth was observed in PL1(1-2), the line with the highest levels of PL1 (Fig. S6a). We observed no difference in titres of DC3000 between NT and any of the transgenic lines (Fig. S6b).

**Figure 2.**
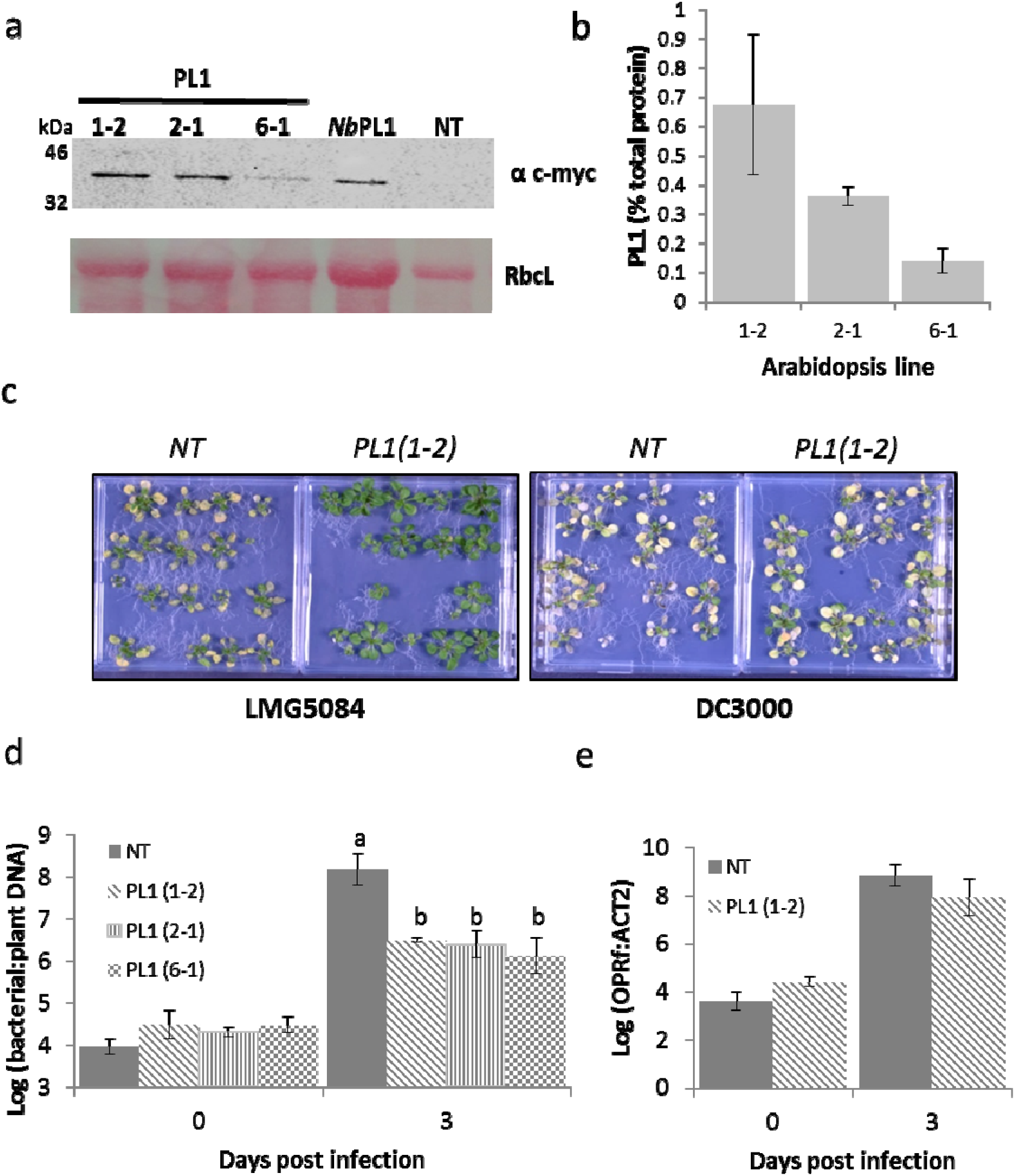
Transgenic Arabidopsis constitutively expressing PL1 attenuates the growth of PL1-susceptible strains, but not PL1 resistant strains. **a**, PL1-can be expressed to high levels in Arabidopsis visualised by western blot of the 3 independent myc-tagged PL1 transgenic lines (1-2, 2-1 and 6-1) in Arabidopsis whole seedlings with the corresponding Ponceau S staining. This data is supplemented with a positive control (*Nb*PL1) from *N. benthamiana* and a non-transgenic Arabidopsis as a negative control (NT). **b**, serial dilutions of whole protein extracts from Arabidopsis seedlings were spotted onto lawns of LMG5084 to estimate the percentage of PL1 activity. Error bars represent the standard deviation of 3 independent replicates. **c**, PL1-expressing seedlings showed qualitative resistance to LMG5084 but not DC3000. 14-day-old Arabidopsis seedlings expressing PL1 and non-transgenic (NT) were flooded with either LMG5084 or DC3000 for 1 minute and symptoms were left to develop over 3 days. 14-day-old Arabidopsis seedlings expressing PL1, PL1(1-2), PL1(2-1) and PL1(6-1), or non-transgenic (NT) were flooded with either **d**, PL1-expressing seedlings showed quantitative resistance to LMG5084 but not DC3000. Quantitative data was LMG5084 or **e**, DC3000 for 1 minute and samples were collected 0 and 3 days post infection. Genomic DNA was extracted from the leaves 0 and 3 days post infection and the bacterial load was measured by qPCR relative to plant tissue. Error bars represent the standard error of 4 independent replicates. Statistical significance within the same time points was revealed using a one-way ANOVA and post hoc Tukey’s T-test. Letters denote statically significant groups (p< 0.05).

As with *N. benthamiana*, titres of bacteria recovered immediately after inoculating PL1-expressing lines with LMG5084 and were lower than expected (Fig. S4). We therefore measured bacterial growth by quantifying DNA levels (Fig. S7). The experiments were carried out using 14-day-old seedlings grown on agar plates because the disease phenotypes are more pronounced on younger plants (25, 26). We observed striking differences in symptom severity between NT and all three PL1 transgenic lines. By 3 dpi, NT seedlings infected with either LMG5084 or DC3000 exhibited severe disease symptoms with most of the seedlings dead or dying. In contrast, for all three PL1-expressing lines nearly all the seedlings infected with LMG5084 appeared green and healthy (Fig. 2c, Fig. S8), whereas those infected with DC3000 showed severe symptoms similar to NT plants (Fig. 2c, Fig. S9). To quantify disease resistance, NT Arabidopsis and the 3 transgenic lines were infected by flooding plates with a suspension of bacteria and samples were taken 0 and 3 dpi. At 3 dpi, quantities of LMG5084 DNA in the PL1 expressing lines were 1.5 -log units and significantly lower (p values for PL1(1-2), PL1(2-1) and PL1(6-1) of 0.002; 0.006; 0.006, respectively) than in the NT control (Fig. 2d). Bacterial DNA levels in PL1(1-2) and NT seedlings infected with the PL1-insensitive line DC3000 were identical (Fig. 2e). In conclusion, PL1 was able to provide qualitative and quantitative disease resistance against the PL1-sensitive strain LMG 5084 but not the insensitive strain DC3000.

### PL1-mediated resistance is not specific to *P. syringae* pv. *syringae* LMG5084

To demonstrate that *in planta* resistance mediated by PL1 expression is not specific to a single strain of *Ps* we tested two additional PL1-susceptible pathovars. We first established which of the remaining 9 PL1 sensitive strains could establish a compatible infection of Arabidopsis by flood-infecting NT seedlings and screening for characteristic *Ps* disease symptoms. We identified *Ps* pv. *syringae* LMG5082 and pv. *lachrymans* LMG5456 as suitable candidates. In flood infections, we observed greatly reduced symptom severity in PL1-transgenic Arabidopsis compared to NT plants with both strains of *Ps* (Fig. 3 a,c; Fig. S10 and S11). Compared to NT plants, bacterial DNA levels in transgenic lines were 0.8-log units lower for LMG5082 and 1.3-log units lower for LMG5456 (Fig. 3 b,d). Therefore, PL1-mediated resistance is not confined to a single strain (LMG5084) nor to *Ps*. pv. *syringae* pathovars.

**Figure 3.**
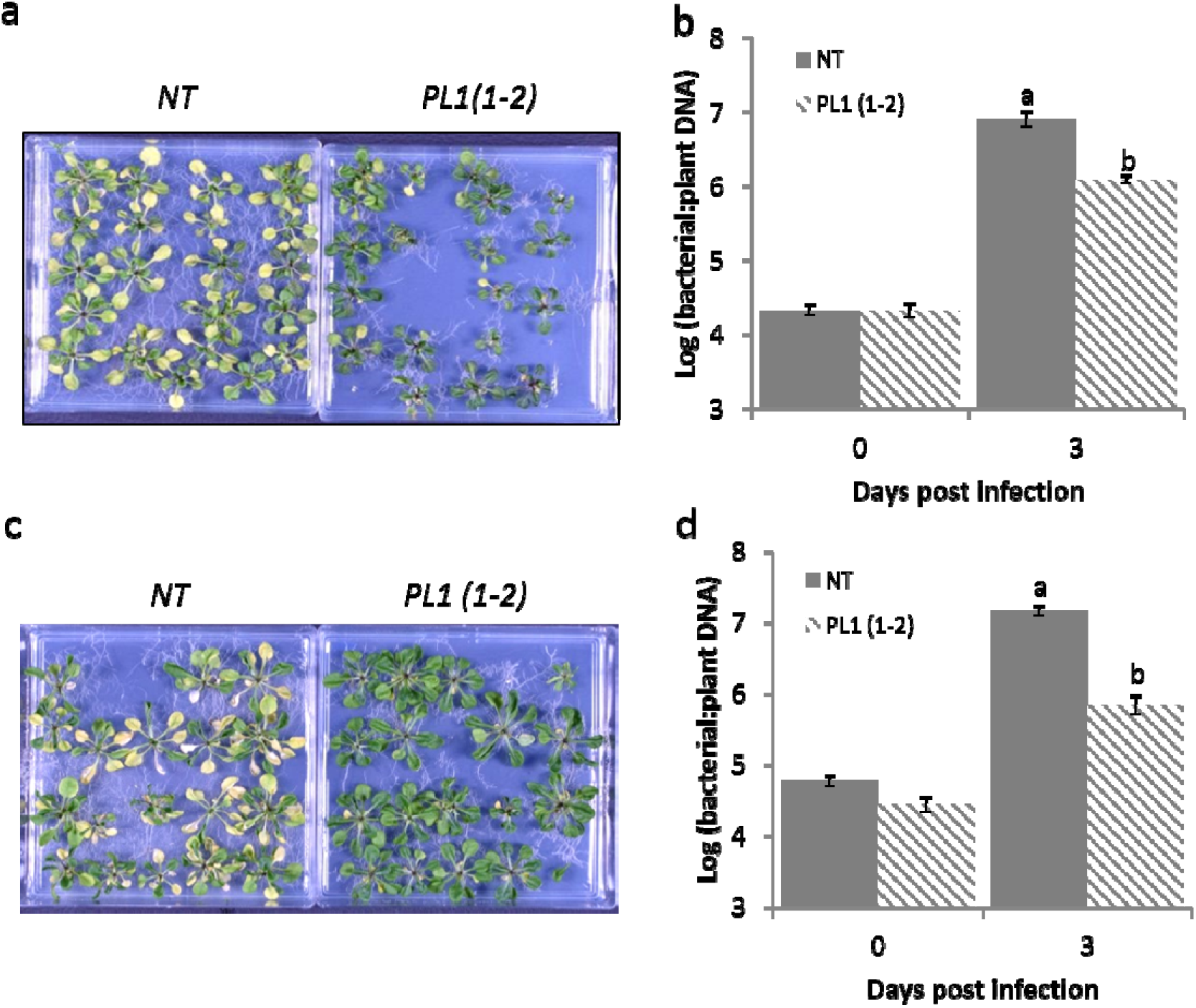
PL1-mediated resistance in Arabidopsis is not specific to LMG5084. Transgenic lines show qualitative resistance to both PL1-sensitive strains. 14-day-old Arabidopsis seedlings NT and PL1(1-2) were flooded with either **a**, LMG5082 or **c**, LMG5456 for 1 minute and symptoms were left to develop over 3 days. The transgenic lines show quantitative resistance to both PL1-sensitive strains. 14-day-old Arabidopsis seedlings WT and 1-2 were flooded with either **b**, LMG5082 or **d**, LMG5456 for 1 minute and samples were collected 0 and 3 dpi. Bacterial DNA was extracted and bacterial load was assessed by measuring the relative amount of bacterial DNA relative to plant DNA by qPCR. Error bars represent standard error of 4 independent replicates. Statistical significance within the same time points was revealed using a one-way ANOVA and post hoc Tukey’s T-test. Letters denote statically significant groups (p< 0.05).

### *P. syringae* mutants that are insensitive to PL1 lack LPS and show reduced virulence in PL-1 transgenic Arabidopsis

Due to the high levels of PL1 being produced *in planta* we predicted that this would put a strong evolutionary pressure on *Ps* to become insensitive to PL1. Previous work has shown that LPS constitutes the primary receptor for the lectin-like bacteriocins and mutations in the LPS synthesis machinery can cause resistance to this class of protein antibiotics (15–17). To begin to assess the robustness of protection to *Ps* provided by *in planta* production of PL1, we selected PL1-insensitive strains by growing LMG5084 in liquid culture in rich media supplemented with 10 µM PL1. Surviving colonies were sub-cultured and tested for their capacity to induce infection in NT and PL1 transgenic Arabidopsis. Eight independent *Ps* mutants that were highly tolerant to PL1, displaying only hazy zones of clearing at >10 µM PL1 in a spot test (Table S2), were selected and whole genome sequencing was performed to identify any mutations that might be responsible for PL1 tolerance. All eight PL1-tolerant strains carried mutations in genes encoding enzymes reported to be involved in LPS biosynthesis (Table S2). To confirm defects in LPS production, we purified and analysed LPS isolated from LMG5084 and the 8 PL1-tolerant strains. Analysis by SDS-PAGE showed that all the mutants lack the O-antigen produced by WT LMG5084 (Figure 4a). In addition, all mutants showed defects in motility as measured in swimming assays and also showed increased sensitivity to reactive oxygen species as determined by exposure to 1% H_2_O_2_ (Figure 4b,c; Table S3 and S4). Interestingly, when NT Arabidopsis were inoculated with the PL1-insensitive strains we did not observe any significant reduction in virulence (based on symptom severity) compared to WT LMG5084. However, transgenic PL1-producing Arabidopsis retained resistance to all PL1-insensitive strains (Figure 4d; Figure S12).

**Figure 4.**
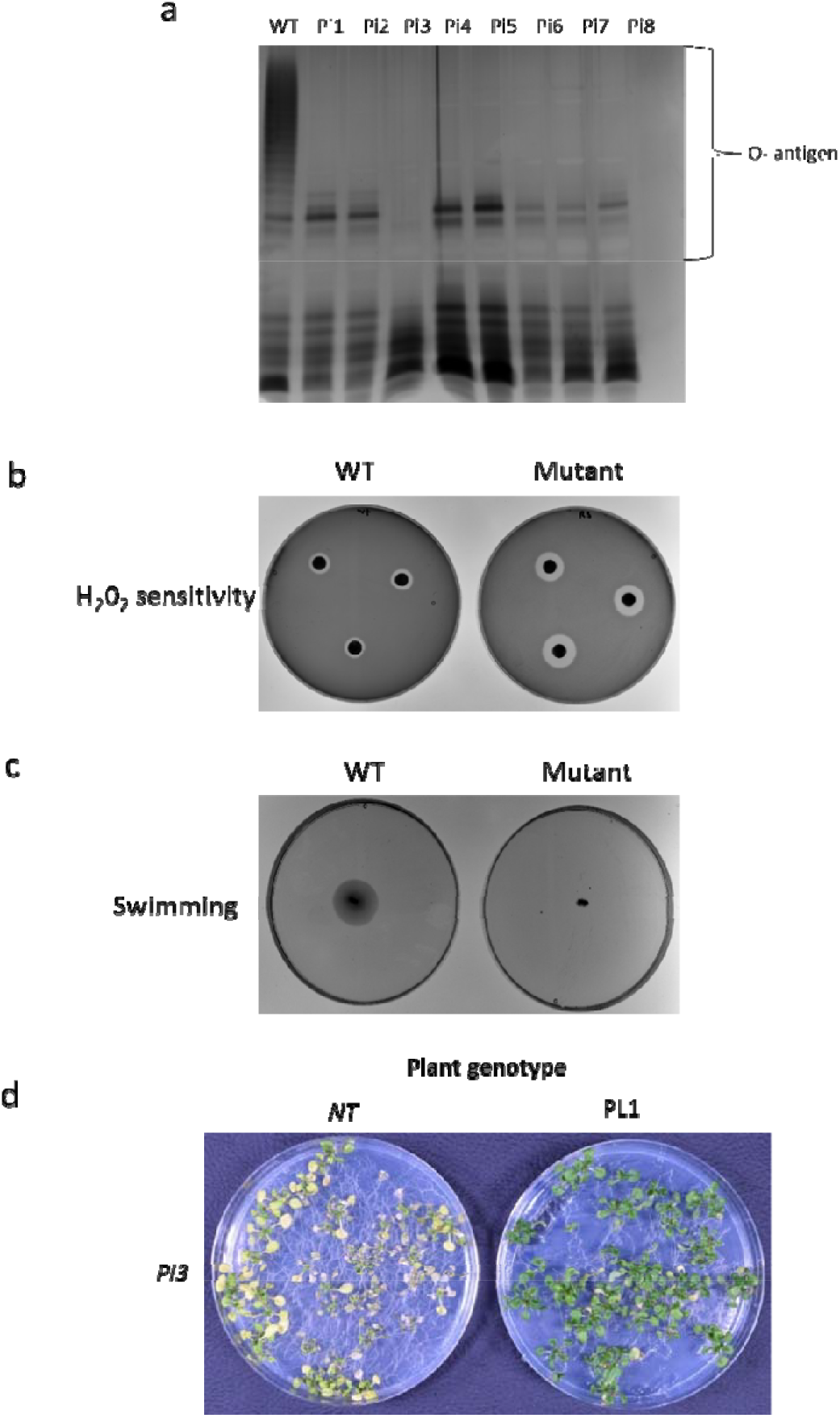
*P. syringae* mutants with PL1-tolerence have mutations in their LPS synthesis genes and cannot overcome PL1-mediated resistance in Arabidopsis. **a**, LPS extracted from PL1-tolerant *Ps* lacks O-antigen. LPS was visualised by SDS PAGE and silver staining. **b**, PL1-tolerant *Ps* show increased sensitivity to hydrogen peroxide. Whatman paper soaked with 1 % H_2_0_2_ was placed onto lawns of wild-type (WT) and mutant *Ps* and incubated overnight. **c**, PL1-tolerant *Ps* display defective motility. Bacterial swimming of WT and mutant *Ps* was assessed in 0.3% hrp-derepressing media and incubated for 4 days. **d**, PL1-tolerant *Ps* are virulent in non-transgenic plants, but not in PL1-transgenics. 14-day-old non-transgenic (NT) or PL1 transgenic (PL1) seedlings were flood inoculated with the mutant bacteria and symptoms were observed after 3 days. Experiments were repeated 3 times showing similar results.

## Discussion

We have established a proof-of-principle for bacteriocin-mediated resistance against a key genus of plant pathogenic bacteria in two different model plant species. We are therefore optimistic that the concept of bacteriocin-mediated crop protection is viable. Encouragingly where bacteriocins have previously been assessed for safety by the US FDA they have been classified as “Generally Regarded as Safe” (18). Furthermore, bacteriocins are narrow-spectrum antimicrobial agents and they should therefore selectively target only specific plant pathogenic bacterial species and not affect the many commensal/mutually beneficial bacteria that persist in the plant rhizosphere, however, this requires further investigation (27). The combination of highly specific target range and negligible impact on benign species has been crucial for the extensive adoption of BT-insecticidal GM crops and we see parallels with the use of bacteriocins for protection against bacterial infections.

To future proof the use of bacteriocins in agriculture we aim to use heterologous cocktails of bacteriocin proteins to overcome the development of bacteriocin insensitivity and to ensure the eradication of plant pathogens (28). We have shown that lectin-like bacteriocins represent promising candidates for transgenic expression. However, genome mining has identified different classifications of bacteriocins in plant pathogenic bacteria like tailocins, colicin-M like bacteriocins and nuclease bacteriocins (29, 30). An important consideration of expressing bacteriocins *in planta* is the effect on plant growth and development. This has not been addressed experimentally, but will need to be addressed in the future. From initial observations there is no phenotypic difference between the transgenic plants and the wild-type. The most vital observation that will be needed to look at is whether there is a yield penalty associated with the expression of PL1 and other lectin-like bacteriocins.

Another consideration is the selection of bacterial populations that develop insensitivity to a bacteriocin, however, this can come with a fitness cost. For example, non-pathogenic strains of Agrobacterium that express agrocin 84 were able to suppress the formation of crown gall by pathogenic strains in the field (31, 32). Our results suggest that for the PL1 tolerant mutants tested here, which were isolated *in vitro*, the levels of PL1 exposure *in planta* in the transgenic lines remained sufficient to offer robust resistance to infection, although these strains remained virulent in non-transgenic plants.

Bacteriocin production is not exclusive to *Pseudomonas* and the strategy is in principle applicable to a wide variety of important phytopathogens e.g. *Xanthomonas spp.* (greening and blight of rice and banana)(33, 34); *Ralstonia solanacearum* (potato brown rot and bacterial wilt of tomato)(35) and *Pectobacterium and Dickeya spp.* (potato soft rot and black leg) (36–39).

We propose that bacteriocin-mediated resistance in plants represents a technology that can be utilised to control bacterial pathogens in the field. Critically, plant-bacterial ecosystems are dynamic and complex, therefore, we expect that their great genomic diversity will promote bacteriocin evolution and hence a very large exploitable resource for applications.

## MATERIALS AND METHODS

### Bacterial strains

*Ps* isolates (Table S1) were obtained from the National Collection of Plant Pathogenic Bacteria and the Belgian Coordinated Collections of Microorganisms. *Ps* strains were cultured in Kings broth B (KB) media, 20 g L^−1^ Peptone, 1.5 g L^−1^ K_2_PO_4_, 1.5 g L^−1^ MgSO_4_, 10 mL L^−1^ glycerol (pH 7.5).

### Motility experiments

For swimming an O/N liquid culture of *Ps* was stabbed into 0.3 % hrp-derepressing minimal media (10 mM sucrose, 50 mM Potassium phosphate buffer, 7.6 mM (NH_4_)_2_SO_4_, 1.7 mM MgCl_2_, 1.7 mM NaCl, pH5.7) agar and incubated for 4 days (40, 41).

### LPS extraction

A pellet from 2 mL of a culture with OD600 = 1 was washed in MgCl_2_ (to remove any trailing media). LPS was then extracted using an LPS extraction kit (iNtRON Biotechnology, Gyeonggi, South Korea). The LPS pellet was resuspended in 50 µL of 10 mM Tris pH 8. To ensure complete solubilisation of the LPS pellet, the sample was boiled at 95 °C for 2 mins and further treated with 3 µg µL^−1^ of proteinase K at 50°C for 30 minutes to obtain a high purity of LPS from bacterial cells.

### Plant growth conditions

*N. benthamiana* plants were grown using long day conditions, 16 hours light/8 hours dark (26/18°C) at 60% humidity. Arabidopsis plants were grown in short day conditions at 80 µmol m^−2^ sec^−1^ with 9 hours light/15 hours dark (22/18°C) conditions at 60/70% humidity.

### Gene cloning

For protein expression in *E. coli*, the sequence encoding PL1 (with no stop codon) was amplified using standard PCR reactions with a high fidelity Phusion Taq polymerase enzyme (New England Biolabs, Hitchin, UK) and appropriate templates, followed by cloning into *NdeI-XhoI* sites in the pET21 vector (16). For constitutive transgene-mediated expression *in planta*, the PL1 coding sequence was fused to an N-terminal 4xMyc tag and cloned into the *kpnI* site of pJO530, a derivative of pBIN19 (42). Furthermore, a GFP vector generated by Cecchini *et al.* was used (42). These plasmids are denoted as pJOPL1 and GFP, respectively. Plasmids used in this study were linearised by digestion using the appropriate restriction enzymes (New England Biolabs, Hitchin, UK). All DNA constructs were verified by sequencing (Source Bioscience).

### Expression and purification of PL1

Expression and purification were carried out according to McCaughey *et al* (16). Briefly, the pET21 plasmid containing PL1 was transformed into BL21 DE3 pLyS cells (Agilent, Edinburgh, UK). PL1 expression was induced at mid-log phase by supplementing the media with 0.3 mM isopropyl β-D-1-thiogalactopyranoside (IPTG), and the cells were grown at 22°C for 20 hours and harvested by centrifugation. The cells were lysed using an MSE Soniprep 150 (Wolf Laboratories, York, UK) and the cell-free lysate was applied to a 5 mL HisTrap HP column (GE Healthcare, Amersham, UK) and PL1 was eluted using a 5–500 mM imidazole gradient. The remaining contaminants were removed by gel filtration chromatography on a Superdex S75 26/600 column (GE Healthcare, Amersham, UK). PL1 was concentrated using a centrifugal concentrator (Vivaspin 20, Epsom, UK) with a 5 kDa molecular weight cut off and stored at −80°C.

### Soft agar overlay susceptibility assays

Soft agar overlay spot assays were performed using the method of Fyfe *et al* (43). Fifty microliters of test strain culture at mid-log was inoculated in 0.8 % soft agar and poured over a KB agar plate, as appropriate. 5 µl of undiluted and serially diluted bacteriocin solution/ plant protein extract was spotted onto the plates and incubated for 20 hours at 28°C, after which plates were inspected for zones of bacterial growth inhibition.

### Transgene expression in *planta*

Transgene expression of the T-DNA constructs *in planta* was achieved by transforming into *Agrobacterium tumefaciens* (strain GV3101). In *N. benthamiana*, leaves were infiltrated with GV3101 containing the appropriate vector (44). For Arabidopsis transformations plants were floral dipped according to Zhang *et al* (45). PL1 expression was detected using western blots. Protein was extracted by macerating frozen leaf tissue in 20 mM Tris-HCl, 200 mM NaCl pH 7.5 supplemented with cOmplete™, EDTA-free Protease Inhibitor Cocktail (Roche, West Sussex, UK) and the protein concentration of the supernatant was determined by Bradford assay (Biorad, Perth, UK). Ten micrograms of protein extract was separated onto 16% SDS-PAGE, transferred onto PVDF membrane and proteins were detected using an anti-c-myc monoclonal antibody (sc-40, Santa Cruz Biotech, Texas, USA) and anti-mouse HRP conjugate (W4021, Promega, Southampton, UK) antibodies.

### Infection assays *in planta*

For infection studies in *N. benthamiana* 1 × 10^5^ Cfu mL^−1^ of *Ps* was syringe-infiltrated into selected leaves of 4-6-week old plants according to Hann and Rathjen (46). For Arabidopsis a suspension of 1 × 10^8^ Cfu mL^−1^ of *Ps* supplemented with 0.025% Silwett L-77 (Lehle Seeds, Texas, USA) was sprayed onto the leaves of 6-week old plants until they were visibly wet (25). For flood inoculation of Arabidopsis, agar plates containing 14-day-old seedlings were flooded for 1 min with a bacterial suspension comprising 1 × 10^6^ Cfu mL^−1^ of bacteria supplemented with 0.025% Silwett L-77 (Lehle Seeds, Texas, USA)^20^. Plates containing transgenic seedlings were supplemented with 15 µg mL^−1^ of hygromycin B (Sigma Aldrich, Gillingham, UK).

### Assay of bacterial titres using qPCR

DNA was extracted from infected *N. benthamiana* leaves using a DNAzol kit (Thermo Fisher Scientific, Paisley, UK) according to the manufacturer’s protocol. For DNA extractions from infected Arabidopsis seedlings, plant tissue was frozen in liquid nitrogen and DNA extracted using FastDNA™ SPIN Kit for Soil (MP Biomedicals, California, USA). Bacterial and plant DNA levels were quantified using qPCR essentially as described by Love *et al* (2007) (47). qPCR was performed in an Applied Biosystems StepOnePlus Real-Time PCR System (Thermo Fisher Scientific, Paisley, UK, using Fast SYBR™ Master Mix (Thermo Fisher Scientific, Paisley, UK) and 0.16 μM of primers). Bacterial DNA levels *in planta* were determined using primers for the *Ps OPRF* gene(24) and normalised against the *ACT2* (*AT3G18780*) gene in Arabidopsis(47) or the 18S rRNA genes in *N. benthamiana.* List of primers are shown in Table S5.

### Isolation and whole genome sequencing and analysis of PL1-insensitive *Ps* strains

Bacteria from a 50 µL overnight culture were pelleted at 3,000 × g for 10 minutes and re-suspended in 1 mL of 10 µM PL1 and incubated for 4 hours and plated out on KB plates and incubated overnight. DNA was extracted from wild type *Ps* LMG5084 and its PL1-insensitive mutants using the GenElute Bacterial Genomic DNA Kit (Sigma-Aldrich). Libraries were prepared with the NEBNext Ultra II library kit according to the manufacturer’s instructions and sequenced on the Illumina HiSeq X platform to obtain 150 bp paired end reads with an average depth of 40-fold. Raw sequences from this study have been deposited in the European Nucleotide Archive (ENA) under the accession numbers detailed in Table S6. *Ps* LMG5084 wild type was assembled using Velvet v 1.2 with multiple assemblies generated using VelvetOptimser v 2.2.5 (48–50). The assembly with the best N_50_ was subjected to assembly improvement. Contigs were ordered using Mauve v 2.4.0, scaffolded with SPPACE v 2.0-1 and gaps filled with GapFiller v 1.11-1 (51–53). The assembly was then annotated using Prokka v 1.5 and Roary v 3.11.2 based on the reference *Ps* B728a accession no. NC_007005.1 (54, 55). This reference genome was chosen according to the top species hit from Kraken (56). For *Ps* LMG5084 PL1-insensitive mutants, sequence reads were mapped to *Ps* LMG5084 wild type using BWA v 0.7.17. SNPs were called and filtered using SAMtools mpileup and BCFtools v 0.1.19 (57). Variant calls were then filtered and retained if the depth was greater than 5, quality greater than 50, mapping quality greater than 20 and the depth of reference (forward and reverse) reads were greater than the depth of alternative reads. Variant effects were predicted using SnpEff v 4.3T (58).

### Statistical analysis

Statistical analysis used in this study was performed with Minitab 17 statistical software using one-way ANOVA followed by Tukey’s multiple comparison test or a Dunnett’s test.

## Acknowledgements

We thank members of the Glasgow Bacteriology and Plant Science groups for discussions and Dr Chantal Keijzer for comments on the manuscript. W.M.R. is supported by the University of Glasgow College of Medical, Veterinary and Life Sciences DTP scholarship and R.W.G was supported by the University of Glasgow Lord Kelvin Adam Smith scholarship. This work was funded by Wellcome Trust grant 201505/Z/16/Z and BBSRC grant BB/L02022X/1.

