## Supplementary information for "Engineering bacteriocin-mediated resistance against plant pathogenic bacteria in plants"

### Supplementary Figures and Tables

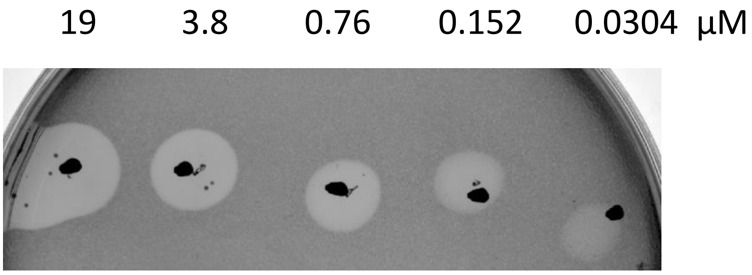


**Figure S1. Representative spot test for sensitivity to PL1.** Serial dilutions of PL1 were spotted onto lawns of the PL1-sensitive strain LMG5084.

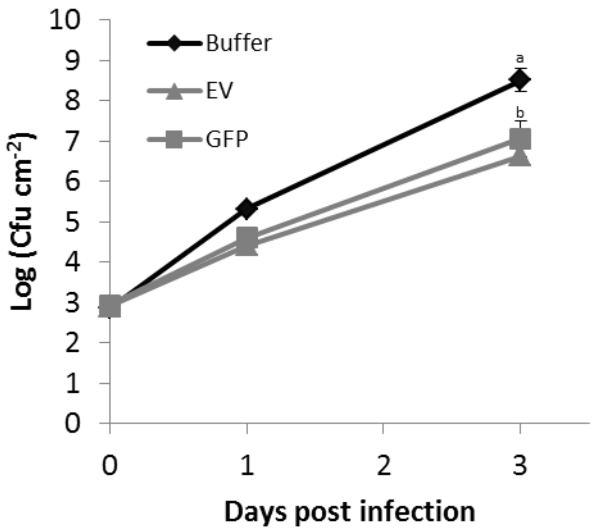


**Figure S2. Bacterial growth in *N. benthamiana* leaves agroinfiltrated with GFP-expressing or empty vector.** *N. benthamiana* leaves were agroinfiltrated with either buffer (diamonds), *Agrobacterium* containing an empty vector or a vector expressing GFP. Plants were then infected with LMG5084 3 days post infiltration, leaf samples were taken 0, 1 and 3 dpi and the bacterial load (Cfu $cm^{-2}$) was measured. Error bars represent standard error of 3 independent replicates. Statistical significance within the same time points was revealed using a 1-way ANOVA post hoc Tukey T-test. Letters denote statically significant groups ($p< 0.05$).

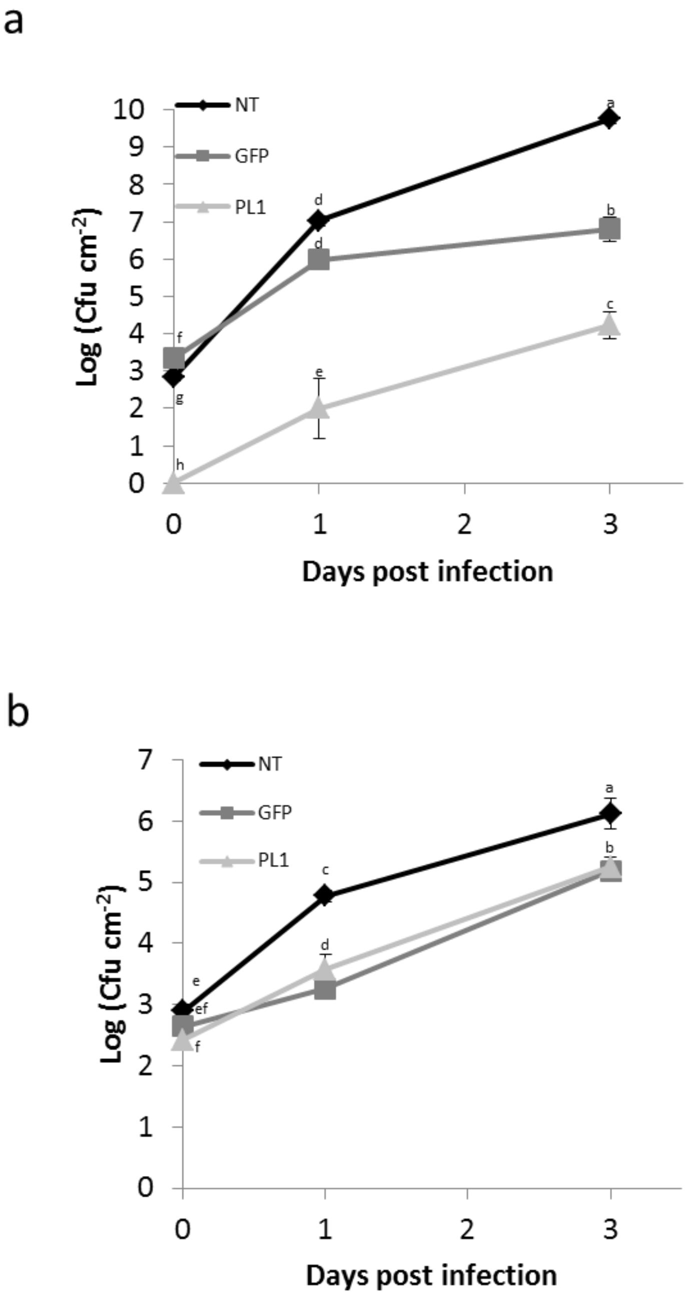


**Figure S3. PL1 expression in *N. benthamiana* attenuates growth of LMG5084 but not DC3000 as (as determined by colony counting).** *N. benthamiana* leaves expressing PL1, GFP or non-agroinfiltrated controls were infected with **a,** LMG5084 or **b,** DC3000**.** Leaf samples were taken 0, 1 and 3 days post infection and the bacterial load (Cfu cm$^{-2}$) was measured. Error bars represent standard error of 3 independent replicates. Statistical significance within the same time points was revealed using a 1-way ANOVA post hoc Tukey T-test. Letters denote statically significant groups ($p < 0.05$).

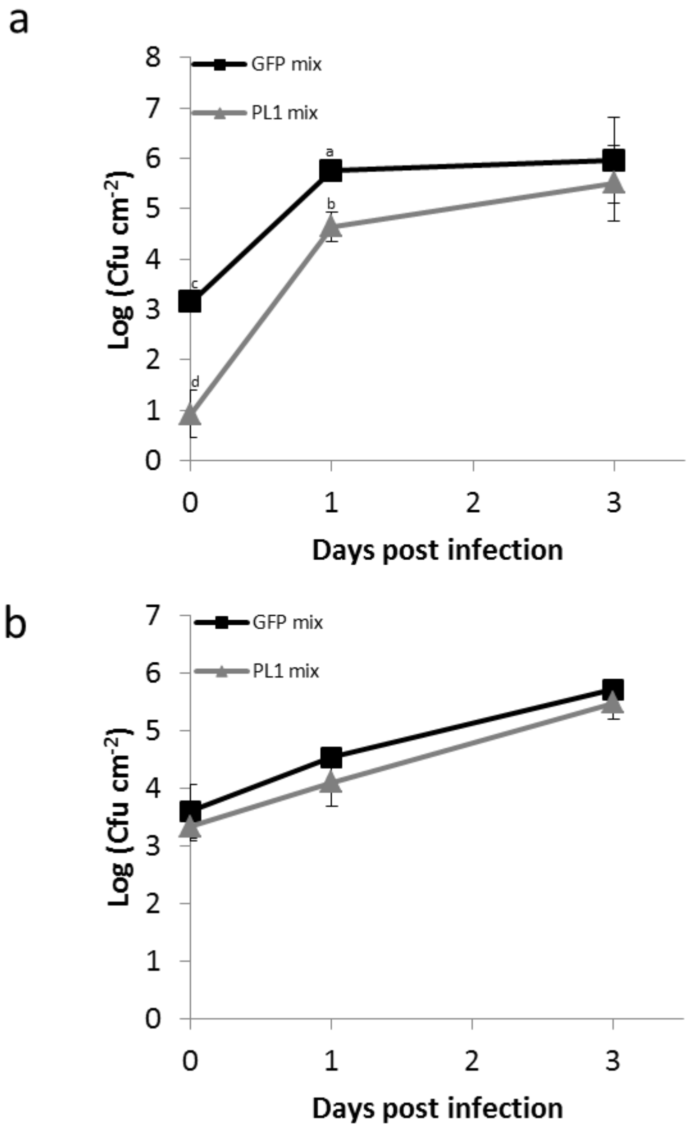


**Figure S4. PL1 expression *in planta* affects bacterial recovery as determined by colony counts.** *N. benthamiana* leaves transiently expressing GFP were syringe-infiltrated with **a,** LMG5084 or **b,** DC3000. Leaf discs were taken 0, 1 and 3 days post infection and mixed with a leaf disc of either a PL1-expressing or an un-infiltrated leaf and, the bacterial loads were (Cfu cm-$^{2}$) measured. Error bars represent standard error of 3 independent replicates. Statistical significance within the same time points was revealed using a 1-way ANOVA post hoc Tukey T-test. Letters denote statically significant groups ($p < 0.05$).

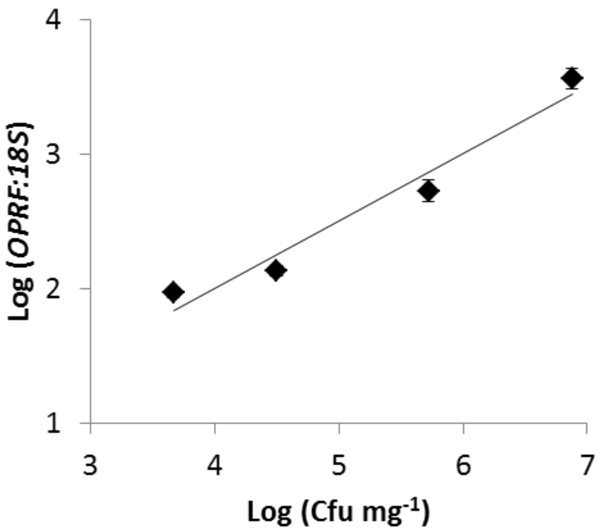


**Figure S5. Bacterial titre in *N. benthamiana* leaves correlates with recovery of bacteria DNA from plant tissue.** Serial dilutions of bacteria were infiltrated into *N. benthamiana* leaves and bacterial DNA was immediately extracted from plant tissue. Levels of bacterial *DNA* relative to *N. benthamiana* DNA were measured using qPCR. Error bars represent standard error of 3 independent replicates.

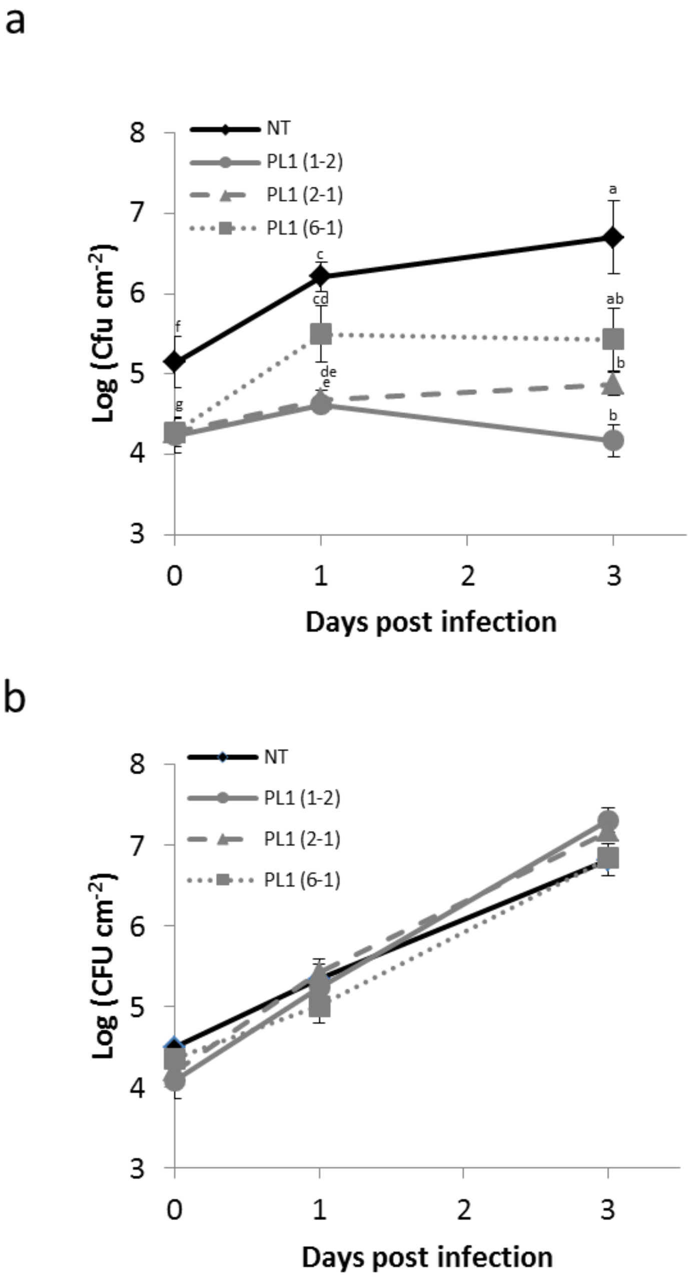


**Figure S6. Arabidopsis expressing PL1 attenuates the growth of LMG5084 but not DC3000 (measured by colony counting).** Three independent PL1 expressing lines, PL1(1-2), PL1(2-1) and PL1(6-1) and a NT control were spray inoculated with 1 x $10^8$ CFU $mL^{-1}$ of either **a,** LMG5084 or **b,** DC3000. Leaf samples were taken 0, 1 and 3 days post infection to measure the bacterial load (Cfu $cm^{-2}$). Error bars represent standard error of 3 independent replicates. Statistical significance within the same time points was revealed using a 1-way ANOVA post hoc Tukey T-test. Letters denote statically significant groups ($p < 0.05$).

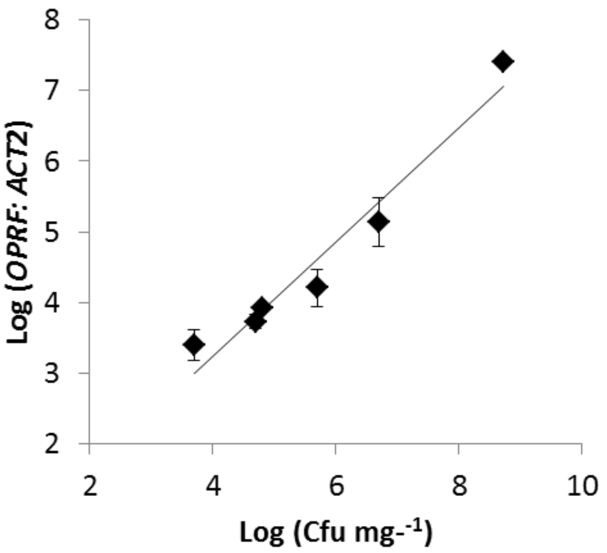


**Figure S7. Bacterial load in Arabidopsis leaves correlates with recovery of bacteria DNA from plant tissue.** Serial dilutions of bacteria were infiltrated into leaves and bacterial DNA was immediately extracted from plant tissue. Levels of bacterial *DNA* relative to Arabidopsis DNA were measured using qPCR. DNA. Error bars represent standard error of 3 independent replicates.

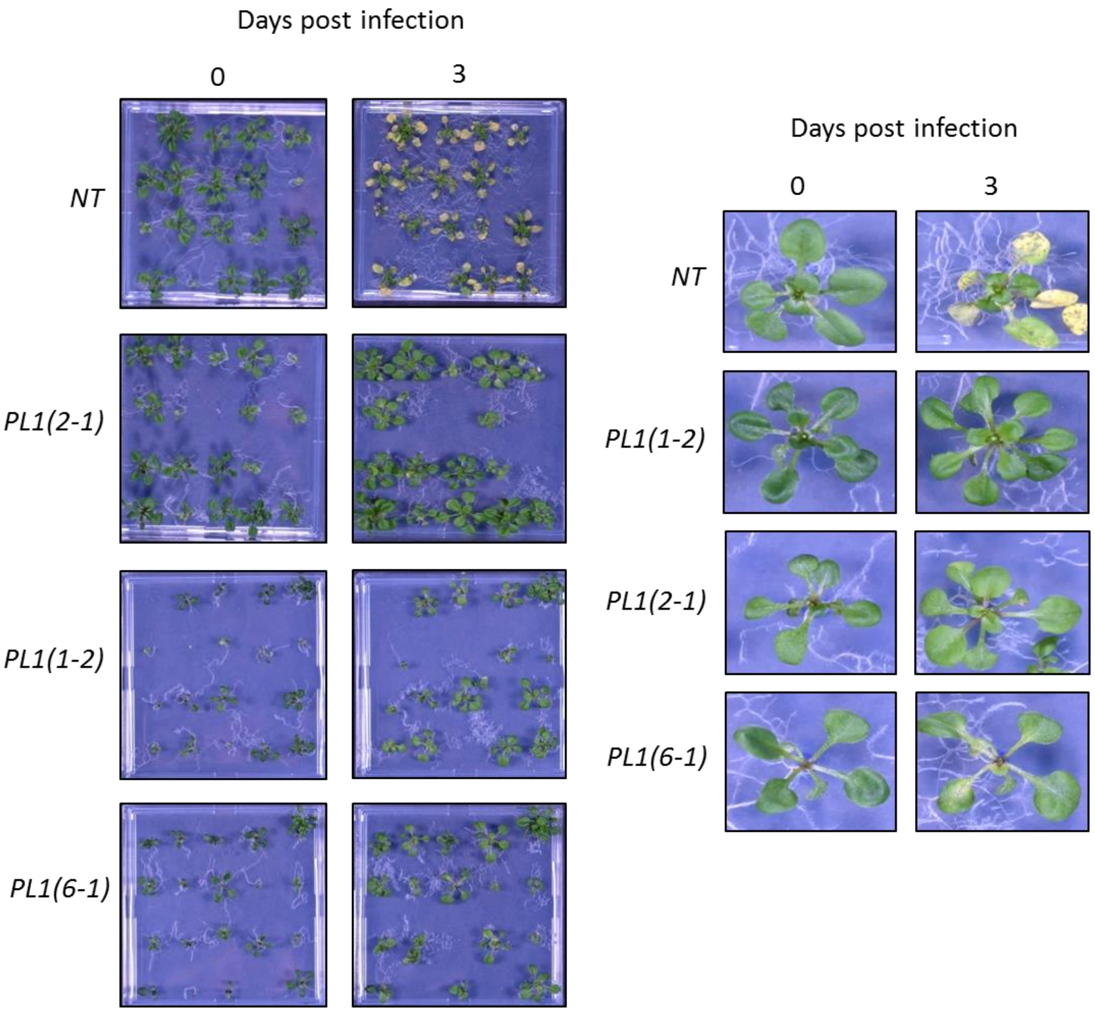


**Figure S8. PL1 expression in Arabidopsis seedlings provides robust disease resistance against LMG5084.** 14-day-old non-transgenic (NT) and 3 independent PL1 transgenic seedlings were flood inoculated with 1 x $10^6$ CFU $mL^{-1}$ of LMG 5084 and symptoms were left to develop other 3 days. Pictures of plates and individual plants were taken 0- and 3-days post infection. Experiments were repeated 3 times with similar results.

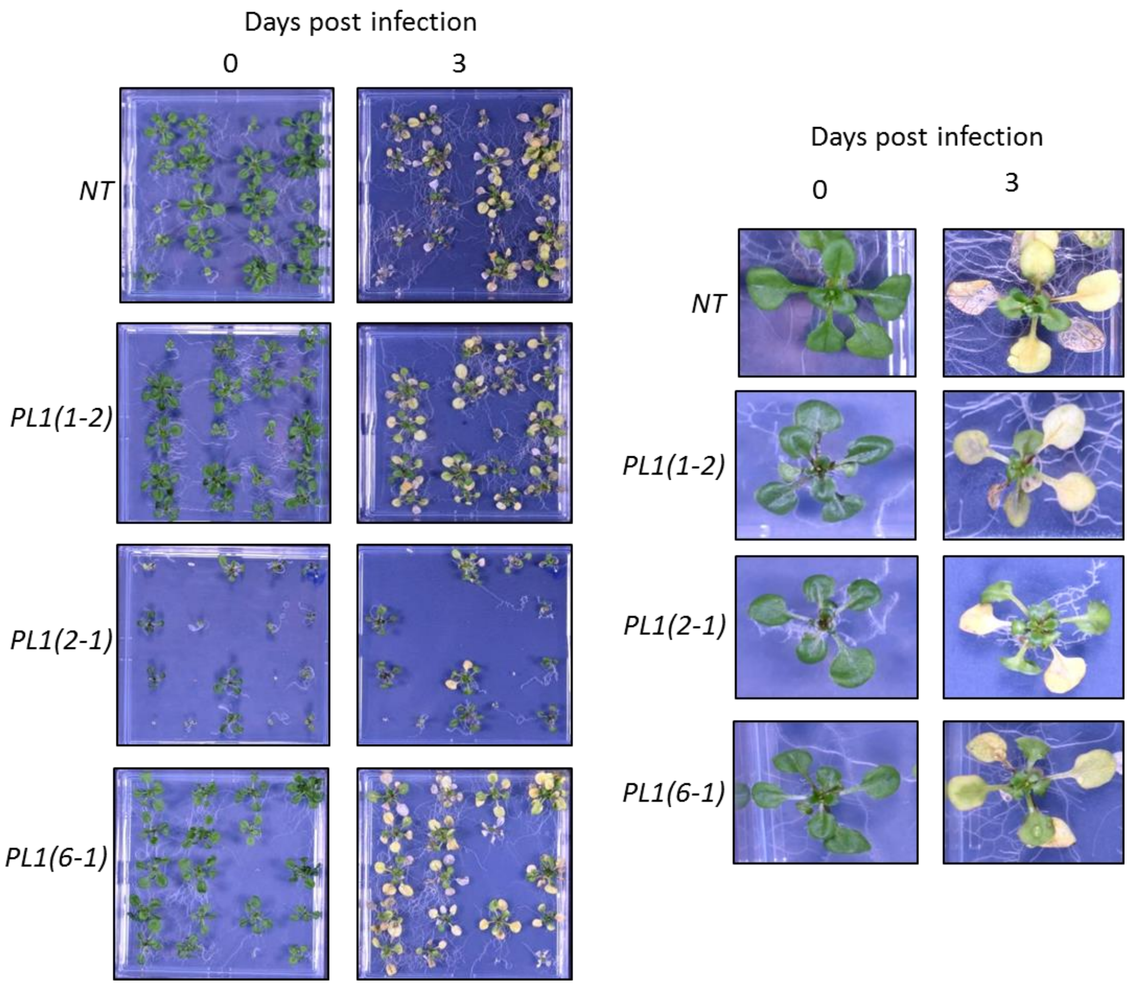


**Figure S9. PL1 expression in Arabidopsis seedlings does not provide robust disease resistance against DC3000.** 14-day-old non-transgenic (NT) and 3 independent PL1 transgenic seedlings were flood inoculated with 1 x $10^6$ CFU $mL^{-1}$ of DC3000 and symptoms were left to develop other 3 days. Pictures of plates and individual plants were taken 0- and 3-days post infection. Experiments were repeated 3 times with similar results.

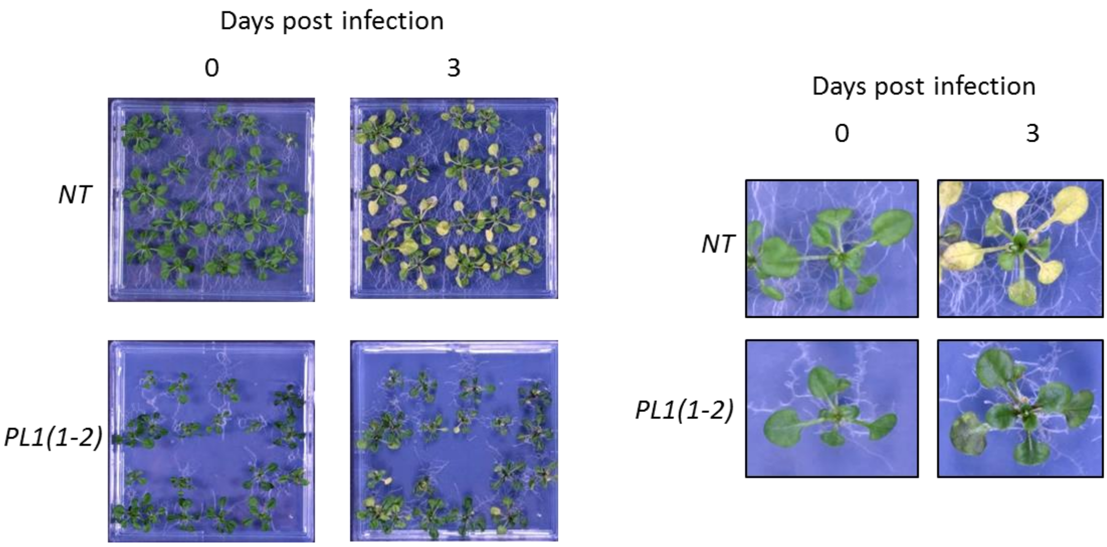


**Figure S10. PL1 expression provides robust disease resistance against LMG5082.** 14-day-old non-transgenic (NT) and transgenic seedlings (PL1(1-2)) were flood inoculated with 1 x $10^6$ CFU $mL^{-1}$ of LMG 5082 and symptoms were left to develop other 3 days. Pictures of plates and individual plants were taken 0- and 3-days post infection. Experiments were repeated 3 times with similar results.

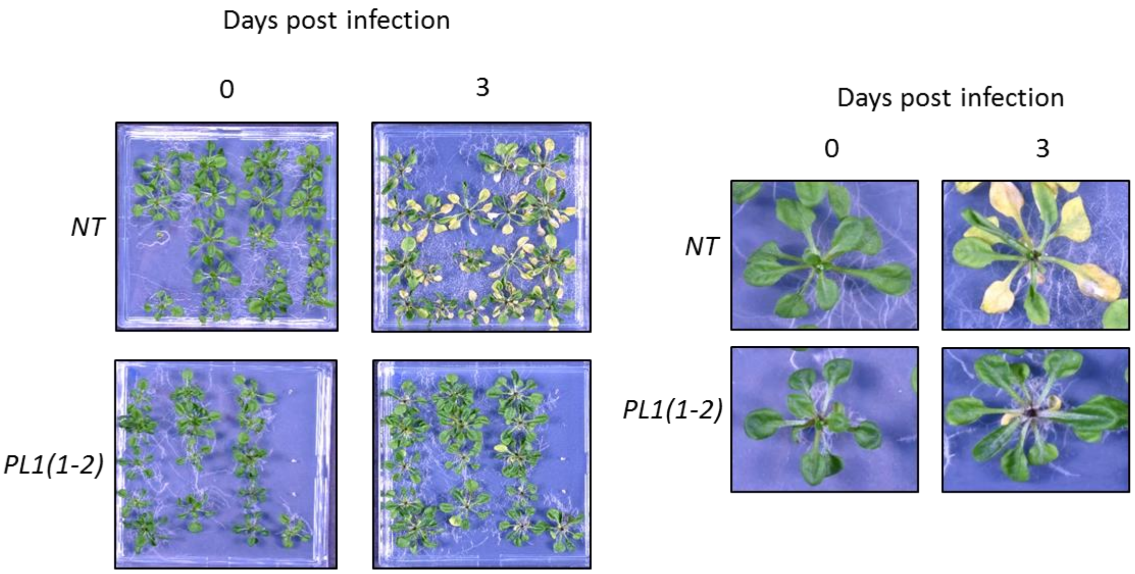


**Figure S11. PL1 expression provides robust disease resistance against LMG5456.** 14-day-old non-transgenic (NT) and transgenic seedlings (PL1(1-2)) were flood inoculated with 1 x $10^6$ CFU $mL^{-1}$ of LMG 5456 and symptoms were left to develop other 3 days. Pictures of plates and individual plants were taken 0- and 3-days post infection. Experiments were repeated 3 times with similar results.

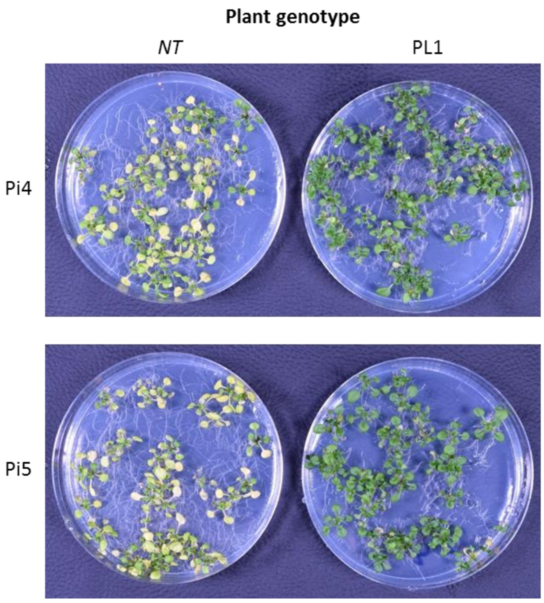


**Figure S12. PL1 insensitive strains of LMG5084 are less virulent in transgenic plants expressing PL1.** 14-day-old non-transgenic (NT) and transgenic seedlings (PL1(1-2)) were flood inoculated with 1 x $10^6$ CFU $mL^{-1}$ of PL1-insensitive *Ps* strains and symptoms were left to develop other 3 days. Pictures of plates and individual plants were taken 0- and 3-days post infection. Experiments were repeated 3 times with similar results.

| Pathovar | Strain ID | Sensitive? | MIC (nM) | Origin | Host |
|---|---|---|---|---|---|
| ***actinidae*** | NCPPB 3738 | Yes | 125 | Japan | *Actinidiae delici* |
| ***actinidae*** | NCPPB 3739 | Yes | 125 | Japan | *Actinidiae delici* |
| ***ciccaronei*** | NCPPB 2355 | No | - | Italy | *Ceratonia siliqua* |
| ***coronafaciens*** | LMG 5060 | No | - | UK | *Avena sativa* |
| ***glycinea*** | NCPPB 2070 | Yes | 1074.2 | USA | *Glycine max* |
| ***glycinea*** | NCPPB 1245 | Yes | 1074.2 | Canada | *Glycine max* |
| ***glycinea*** | NCPPB 2895 | No | - | Australia | *Glycine max* |
| ***glycinea*** | NCPPB 3643 | No | - | Brazil | *Glycine max* |
| ***lachrymans*** | LMG 5456 | Yes | 22.9 | UK | *Cucumis sativus* |
| ***maculicola*** | LMG 2208 | No | - | UK | *Brassica oleraccea* |
| ***morsprunorum*** | LMG 2222 | Yes | 0.85 | UK | *Prunus avium* |
| ***persicae*** | NCPPB 3687 | No | - | New Zealand | *Prunus salicina* |
| ***persicae*** | NCPPB 2254 | No | - | France | *Prunus salicina* |
| ***savastoni*** | NCPPB 1506 | Yes | 325 | Italy | *Olea europaea* |
| ***savastoni*** | NCPPB 2327 | No | - | Italy | *Olea europaea* |
| ***syringae*** | LMG 5084 | Yes | 5.6 | UK | *Pyrus communis* |
| ***syringae*** | LMG 5082 | Yes | 8.3 | UK | *Zea Mays* |
| ***syringae*** | LMG 1247 | Yes | 1851.85 | UK | *Syringa vulgaris* |
| ***tomato*** | NCPPB 3160 | No | - | UK | *Solanum lycopersicum* |
| ***tomato*** | NCPPB 2563 | No | - | UK | *Solanum lycopersicum* |
| ***tomato*** | NCPPB 1107 | No | - | UK | *Solanum lycopersicum* |
| ***tomato*** | DC3000 | No | - | USA | *Solanum lycopersicum* |

**Table S1. The range of *P. syringae* pathovars that are susceptible to PL1.**

| Mutant strain | PL1 sensitivity | Predicted gene | Position in ORF (bp) | sequence 5’→3’ (WT v mutant) |
|---|---|---|---|---|
| Pi1 | >10 µM[a] | Glycosyltransferase; GenBank: CP005969.1 | 588 | AAGTTCTTTCTGTTCGTATTAC<br>AAGT--------------------------ATTAC |
| Pi2 | >10 µM[a] | wbpL; glycosyltransferase; GenBank: CP005969.1 | 333 | GCCTGGGCTTGT<br>GCCT--GGCTTGT |
| Pi3 | >10 µM[a] | wpbM; nucleoside-diphosphate sugar epimerase; GenBank: CP005969.1 | 285 | CTGCGGGAAACC<br>CTGCGG--AAACC |
| Pi4 | >10 µM[a] | Phosphomannomutase; GenBank: CP005969.1 | 1298 | CCGCGCCATCGGC<br>CCGCG**A**C**G**TCGGC |
| Pi5 | >10 µM[a] | GDP-mannose 4,6-dehydrase; GenBank: CP005969.1 | 763 | CGTGGCCGTGAG<br>CGTG--CCGTGAG |
| Pi6 | >10 µM[a] | Glycosyltransferase; GenBank: CP005969.1 | 647 | TACCCGGTGGTGATCCTCGGTGGCGGGC<br>TACC ----------------------------GGTGGCGGGC |
| Pi7 | >10 µM[a] | wpbM GenBank: CP005969.1 | 618 | GCGATGCAGT<br>GCGAT--CAGT |
| Pi8 | >10 µM[a] | O-antigen ligase-like GenBank: CP005969.1 | 606 | AGACGCGTACTGCACTGGT<br>AGACG------------------ --TGGT |

**Table S2. Mutations linked with PL1-resisistance.**

>10 µM[a] – hazy zones of clearing observed at >10 µM

| **Strain** | Average diameter (mm) | Significance |
|---|---|---|
| WT | 21.00 | n/a |
| Pi1 | 7 | Y |
| Pi2 | 12 | Y |
| Pi3 | 3 | Y |
| Pi4 | 11 | Y |
| Pi5 | 9 | Y |
| Pi6 | 7 | Y |
| Pi7 | 8 | Y |
| Pi8 | 10 | Y |

**Table S3. Swimming of PL1-insensitive strains in HRP-de-repressing minimal media.** Bacterial cultures were inoculated into HRP de-repressing media supplemented with 0.3% agar and incubated at 24 °C for six days. Experiments were repeated 3 times with similar results (n=3). Statistical significance for the PL1 insensitive mutants compared to the WT was assessed by a one-way ANOVA Dunnett’s post hoc test.

| **Strain** | **Diameter (mm)** | **Significance** |
|---|---|---|
| WT | 10 | **n/a** |
| Pi1 | 14 | **Y** |
| Pi2 | 15 | **Y** |
| Pi3 | 13 | **Y** |
| Pi4 | 14 | **Y** |
| Pi5 | 13 | **Y** |
| Pi6 | 12 | **Y** |
| Pi7 | 12 | **Y** |
| Pi8 | 13 | **Y** |

**Table S4. Sensitivity of PL1-insenstitive to 1% hydrogen peroxide.** Whatman paper socked in 1% $H_20_2$ were placed on lawns of bacteria and incubated at 24 °C overnight. Experiments were repeated 3 times with similar results (n=3). Statistical significance for the PL1 insensitive mutants compared to the WT was assessed by a one-way ANOVA Dunnett’s post hoc test.

| Primer name | Sequence 5’ -> 3’ |
|---|---|
| OPRF_F | AACTGAAAAACACCTTGGGC |
| OPRF_R | CCTGGGTTGTTGAAGTGGTA |
| ACT2_F | CTAAGCTCTCAAGATCAAAGGCTT |
| ACT2_R | ACTAAAACGCAAAACGAAAGCGGT |
| 18S_F | ATTGGAGGGCAAGTCTGGTGC |
| 18S_R | GCA GAA GGG ACG AGA CGA C |

**Table S5. qPCR primers used in this study**

| Sanger ID | Sample | ENA accession number |
|---|---|---|
| 4526STDY7070045 | *Ps* 5084 Parent | SAMEA104233059 |
| 4526STDY7070060 | *Ps* 5084 R1 | SAMEA104233065 |
| 4526STDY7070068 | *Ps* 5084 R2 | SAMEA104233068 |
| 4526STDY7070076 | Ps 5084 R3 | SAMEA104233071 |
| 4526STDY7070084 | *Ps* 5084 R4 | SAMEA104233074 |
| 4526STDY7070092 | *Ps* 5084 R5 | SAMEA104233077 |
| 4526STDY7070100 | *Ps* 5084 R6 | SAMEA104233080 |
| 4526STDY7070108 | *Ps* 5084 R7 | SAMEA104233083 |
| 4526STDY7070116 | *Ps* 5084 R8 | SAMEA104233086 |
| 4526STDY7070124 | *Ps* 5084 R9 | SAMEA104233089 |
| 4526STDY7070037 | *Ps* 5084 R10 | SAMEA104233055 |

**Table S6. ENA accession numbers for sequenced samples**
